# Hidden assumptions in nascent RNA sequencing pipelines define reproducibility states

**DOI:** 10.64898/2026.07.13.738089

**Authors:** Xinbo Zhou, Cui Feng, Yixin Zhao

## Abstract

Reproducibility of sequencing analyses is often assumed when identical data are processed with established pipelines, yet outcomes can depend on library assumptions that are not explicit to users. Here we compared commonly used pipelines for nascent RNA sequencing. Across public human PRO-seq datasets, identical inputs produced structured divergence in transcriptional profiles. A diagnostic workflow traced this divergence to interactions among paired-end library design, UMI organization, read trimming and alignment strategy. Similar patterns were observed in independently generated human and pig PRO-seq libraries sharing a dual-end UMI design, including divergence associated with pipeline behavior that could not be altered through user-accessible parameters alone. Beyond PRO-seq, GRO-seq analyses showed that assay-specific library architecture and signal-coordinate conventions could distort positional profiles even without UMI processing. In PRO-cap and re-examined PRO-seq datasets, incomplete UMI metadata either prevented pipeline execution or caused silent signal loss; unreported terminal UMIs were detected in four of five examined PRO-seq datasets. Together, these results define reproducibility states shaped by library design, pipeline assumptions and metadata availability.

## Introduction

Reproducibility is a central expectation in computational analyses of high-throughput sequencing data^1,2^. In principle, identical sequencing datasets processed with established computational pipelines should produce consistent analytical outcomes. In practice, however, different bioinformatics tools applied to the same raw data can yield substantially different results^3–6^. Such divergence complicates the interpretation of sequencing experiments and raises questions about computational pipelines that are often treated as routine components of genomic research^7^.

Divergence between analytical outputs is often attributed to differences in software implementation or parameter choice^8–10^. Beyond these explicit choices, pipelines also encode assumptions about sequencing library structure, read orientation and metadata that may not be fully documented or visible to users. When libraries with different architectures are processed under these assumptions, systematic divergence can arise in ways that are difficult to anticipate or diagnose. Understanding how such hidden assumptions affect analytical outcomes is therefore critical for interpreting reproducibility in sequencing-based studies.

Nascent RNA sequencing assays provide a useful context in which to examine these issues. Methods such as PRO-seq^11^, GRO-seq^12^ and related approaches^13,14^ directly profile engaged RNA polymerases and generate genome-wide maps of transcriptional activity^15^. These assays are widely used to study transcriptional regulation and rapid transcriptional responses across diverse biological contexts^16–21^. Beyond descriptive profiling, nascent RNA sequencing data are increasingly used as inputs for quantitative models of transcriptional regulation, including models of initiation^22,23^, promoter-proximal pausing^24^, elongation^25–27^, and RNA stability^28^. Accurate processing of positional nascent RNA signals is therefore important not only for reproducible genome browser tracks, but also for downstream models that infer biological processes from read distributions.

At the same time, nascent RNA sequencing assays vary substantially in library design and preparation strategy^29,30^. Differences in read configuration, adapter architecture and unique molecular identifier (UMI) usage are common across studies, yet are not always fully described in public metadata^31,32^. When such library structures are processed using standard pipelines, interactions between library design and pipeline assumptions become particularly evident. Nascent RNA sequencing therefore provides an informative system for examining how experimental protocol variation, pipeline assumptions and incomplete metadata jointly shape analytical reproducibility.

Here we use published human PRO-seq datasets as an initial case study to identify structured divergence in transcriptional profiles generated by commonly used pipelines from identical sequencing inputs. Diagnostic analyses trace this divergence to interactions between paired-end library design, UMI processing and pipeline-embedded assumptions, rather than to individual pipeline effects alone. We further show that related divergence patterns persist in independently generated human libraries, across species and across related nascent RNA assays. Together, these analyses define reproducibility states that arise from interactions among library design, pipeline assumptions and metadata availability.

## Results

### A framework for classifying reproducibility states in nascent RNA sequencing analysis

Nascent RNA sequencing technologies are widely used to profile transcriptional activity^12,33,34^, yet systematic divergence can arise when identical datasets are processed with different established pipelines, and such divergence is often difficult to attribute to a single source. To interpret these outcomes, we considered three interacting factors: sequencing library design, pipeline assumptions and metadata availability (**Figure 1A**). Library design includes features such as read configuration, adapter architecture and UMI usage. Pipeline assumptions include preprocessing, read alignment, filtering and signal-generation strategies. Metadata availability refers to whether pipeline-relevant information is accessible to users in a form that enables appropriate parameter specification and interpretation. Together, these factors can influence not only how sequencing reads are processed, but also how transcriptional signal profiles are generated and interpreted downstream.

**Figure 1.**
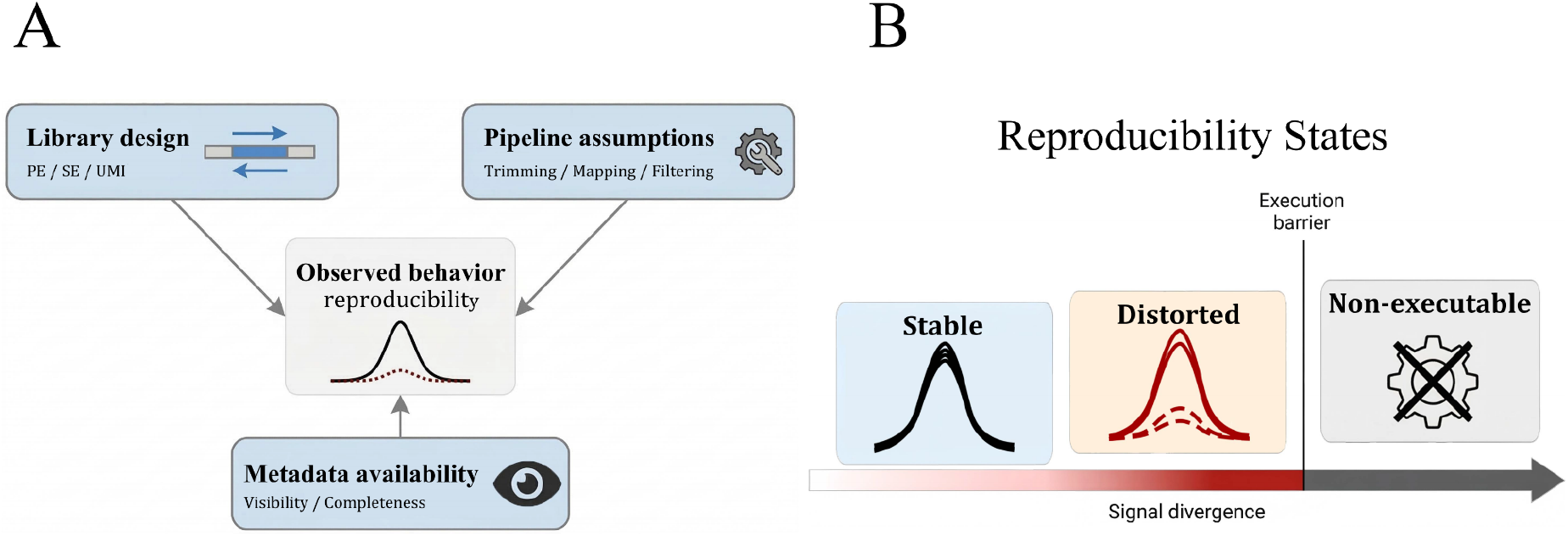
Conceptual framework of reproducibility outcomes and state definitions. (A) Analytical outcomes are influenced by three interacting factors: the design of the sequencing library (library design, including single- or paired-end strategies and UMI usage), the assumptions used in computational pipelines (pipeline assumptions, including read trimming, mapping, and filtering), and the availability of essential experimental information (metadata availability). (B) Reproducibility outcomes can be grouped into three states: the stable state, in which analytical outputs remain consistent across pipelines; the distorted state, in which outputs diverge systematically despite identical inputs; and the non-executable state, in which analysis cannot be completed under available metadata conditions because essential pipeline-relevant information is missing or inaccessible.

We therefore defined three reproducibility states (**Figure 1B**). In the stable state, analytical outputs remain consistent across pipelines and preserve comparable transcriptional signal profiles. In the distorted state, pipelines produce systematically divergent outputs despite operating on identical input data. In the non-executable state, analysis cannot be completed under available metadata conditions because essential pipeline-relevant information is missing or inaccessible. These states provide a practical vocabulary for describing reproducibility outcomes across the analyses below.

### Pipeline-associated divergence from identical PRO-seq datasets

We first asked whether commonly used nascent RNA sequencing pipelines produce consistent transcriptional profiles from identical PRO-seq inputs under a controlled biological context (**Supplementary Figure S1**). To address this question, we curated 18 published human K562 PRO-seq libraries from 10 studies^16,19,28,35–41^ and analyzed each dataset using three widely adopted pipelines: proseq2.0^42^, PEPPRO^43^ and nf-core/nascent^44,45^. We focused on K562 datasets to minimize variation in biological source material while retaining diversity in library architecture, allowing us to distinguish pipeline-associated divergence from differences driven primarily by cell type or experimental context. These untreated samples represented different library designs, including single-end libraries, paired-end libraries without UMIs and paired-end libraries with UMIs (**Supplementary Table S1**). Library design annotations were curated from published manuscripts, supplementary materials and associated GEO records, reflecting the metadata typically available to users for downstream re-analysis.

Hierarchical clustering based on Pearson correlations of log_2_-transformed reads per million (RPM) signal intensities within ±500 bp of transcription start sites (TSSs) showed that samples grouped primarily by analysis pipeline rather than by dataset origin (**Figure 2A**; see **Methods**). Within each pipeline, sample-to-sample correlations remained high (Pearson’s r > 0.8), whereas correlations across pipelines were consistently lower, often falling below 0.7. Similar patterns were observed when signal quantification was extended to gene body regions (**Supplementary Figure S2A**). These observations indicate that cross-pipeline divergence is structured rather than random.

**Figure 2.**
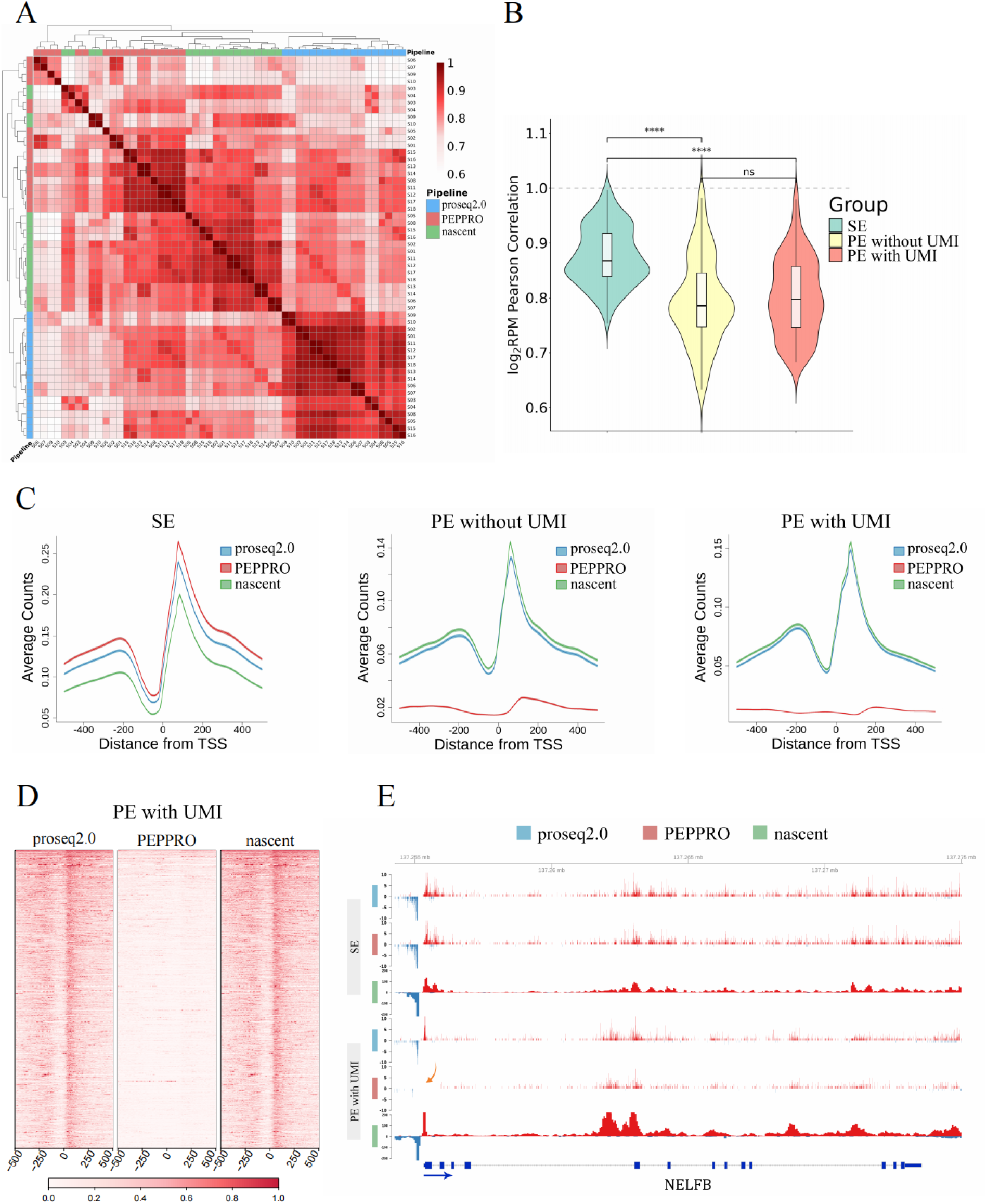
Structured divergence in PRO-seq datasets processed by proseq2.0, nf-core/nascent, and PEPPRO. (A) Clustered heatmap of log_2_RPM-based correlations for transcriptional profiles generated by the three pipelines around transcription start sites (TSSs; ±500 bp). The color gradient reflects correlation magnitude (red, high; white, low), and side color bars indicate the corresponding pipelines. (B) Violin plots showing the distribution of TSS-region correlations across three library design groups (single-end, paired-end without UMIs, and paired-end with UMIs). The y-axis represents Pearson correlations of log_2_RPM signals for pairwise comparisons within each group. Statistical significance was assessed using pairwise Wilcoxon rank-sum tests (****, *p-adj* < 1 ✕ 10^-17^). (C) Metaplots of signal profiles around TSS regions for representative samples from each library design group, including single-end (Sample S11, see **Supplementary Table S1**), paired-end without UMIs (Sample S06), and paired-end with UMIs (Sample S01) libraries. The x-axis indicates distance from the TSS, and the y-axis shows average raw counts. Curves correspond to transcriptional profiles generated by the three pipelines. (D) Gene-level heatmaps of TSS-proximal signal intensities for paired-end libraries containing UMIs (Sample S01). Rows represent TSS-centered windows from the top 10% of genes ranked by mean promoter-proximal signal across pipelines, and columns represent binned positions across the TSS-centered region. Heatmaps are shown separately for proseq2.0, PEPPRO and nf-core/nascent outputs. Signal values were scaled and clipped to 0–1 for visualization and plotted using a common color scale across the three pipelines. (E) Genome browser tracks at a representative locus (NELFB), illustrating local signal profiles for single-end (Sample S11) and paired-end (Sample S01) libraries containing UMIs processed by each pipeline. The blue arrow indicates the direction of NELFB transcription, whereas the pink arrow indicates reduced signal near the TSS region. Red and blue tracks denote signals mapped to the positive and negative strands, respectively. Note that BigWig files generated from proseq2.0 and PEPPRO are single-nucleotide resolution, while nf-core/nascent shows interval-based resolution.

We next examined whether this divergence was associated with library design. Single-end libraries showed relatively consistent transcriptional profiles across pipelines (median Pearson’s r = 0.87; **Figure 2B**). In contrast, paired-end libraries, both with and without UMIs, showed lower cross-pipeline correlations than single-end libraries. Correlations also differed among pipeline comparisons within each library-design group (**Supplementary Figure S2B**). Together, these results associate increased cross-pipeline divergence with paired-end library architecture and its interaction with pipeline behavior.

To assess how these differences manifest as positional transcriptional signals, we examined TSS-centered metaplot profiles (see **Methods**). Representative profiles are shown in **Figure 2C**, with consistent patterns across samples within each library group (**Supplementary Figure S2C**). For single-end libraries, all three pipelines produced broadly similar profiles with comparable enrichment around the TSS, consistent with the reduced divergence observed in correlation-based analyses (**Figure 2B**). By contrast, paired-end libraries showed distinct profiles across pipelines, particularly in promoter-proximal regions downstream of the TSS. Under these library designs, proseq2.0 and nf-core/nascent produced broadly similar profiles, whereas PEPPRO showed reduced promoter-proximal signal intensity (**Figure 2C**).

To determine whether this reduction reflected a broadly distributed pattern rather than an average-profile effect, we visualized TSS-proximal signals across genes with the strongest promoter-proximal signal, ranked by the mean signal across pipelines (see **Methods**). In paired-end libraries with UMIs, proseq2.0 and nf-core/nascent showed coherent promoter-proximal enrichment across individual TSS windows, whereas PEPPRO showed broadly reduced signal intensity (**Figure 2D**; additional library-design groups in **Supplementary Figure S2E**).

Because promoter-proximal and gene-body signals are commonly summarized using metrics such as pausing index in transcriptional analyses, we further examined whether pipeline-associated signal differences propagated to downstream quantitative estimates (see **Methods**). In paired-end libraries with UMIs, PEPPRO showed a lower pausing-index distribution than proseq2.0 and nf-core/nascent, with median log_2_(pausing index) values of 2.98, 3.86 and 3.62, respectively. In contrast, single-end libraries showed more comparable distributions across pipelines (**Supplementary Figure S2F**).

Representative locus-level examples further illustrated these divergence patterns at individual genes. At the NELFB locus, promoter-proximal signal profiles were consistently represented across pipelines in single-end libraries, whereas reduced signal intensity was observed in paired-end libraries containing UMIs when processed with PEPPRO (**Figure 2E**; pink arrow). An additional locus showing a similar pattern is provided in **Supplementary Figure S2D**. Together, these analyses show that divergence from identical PRO-seq inputs is structured, associated with library design and reflected in positional transcriptional signal profiles at multiple levels. We therefore hypothesized that this divergence arose from interactions among specific processing steps rather than from any individual component, and tested this hypothesis using a diagnostic workflow.

### Structured divergence arises from interactions between processing steps

To understand how the structured divergence observed in **Figure 2** arises, we examined major processing steps shared across pipelines, including read trimming, sequence alignment, post-alignment filtering and transcriptional signal generation (**Supplementary Figure S3A**). Rather than treating pipelines as indivisible units, we compared how these steps are combined within each pipeline. For paired-end libraries with UMIs at both read ends, proseq2.0 and nf-core/nascent apply dual-end trimming strategies, whereas PEPPRO uses a 5′-end trimming approach. These strategies differ in how barcodes and UMIs are identified and removed, particularly in the presence of read-through events, leading to distinct read boundaries (**Supplementary Figure S3B**). The pipelines also differ in their alignment and filtering strategies: proseq2.0 and nf-core/nascent use bwa^46,47^ for read mapping, whereas PEPPRO uses bowtie2^48^; post-alignment filtering is applied by default in proseq2.0 and PEPPRO but not in nf-core/nascent. All three pipelines generate genome-wide signal tracks, but differ in signal resolution and coordinate conventions (see **Methods**).

To test whether specific combinations of processing steps could recapitulate the observed divergence, we constructed a diagnostic workflow that systematically varied combinations of trimming strategy, aligner and post-alignment filtering (**Figure 3A**; see **Methods**). This analysis focused on paired-end libraries with UMIs, where interactions between trimming and alignment can be most clearly examined. Clustering based on TSS-proximal signal showed that 5′-end trimming with bowtie2 alignment formed a distinct cluster, whereas the other processing combinations produced broadly similar profiles (**Figure 3B**). This divergence pattern persisted in the absence of MAPQ-based post-alignment filtering (**Supplementary Figure S3C**), indicating that it arose before filtering and reflected an interaction between trimming and alignment strategies.

**Figure 3.**
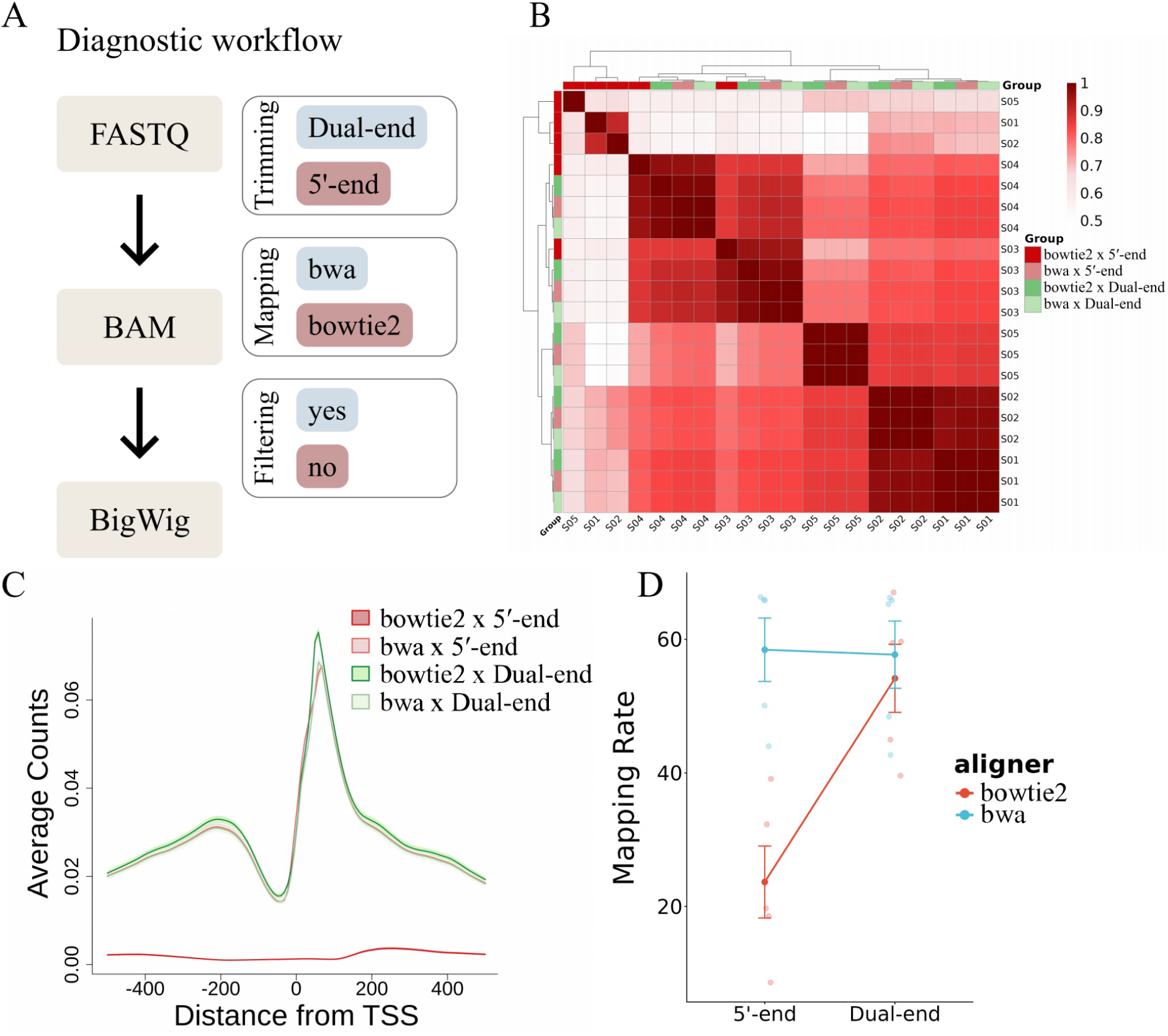
Structured divergence arises from interactions between processing steps in PRO-seq data analysis. (A) Schematic overview of the diagnostic workflow constructed from different trimming strategies, aligners and post-alignment filtering conditions to examine interactions between processing steps. (B) Clustered heatmap of log_2_RPM-based correlations around TSS (±500 bp) for processing combinations generated by the diagnostic workflow. One processing combination (bowtie2 with 5′-end trimming) forms a distinct cluster relative to the other combinations. (C) Metaplots of signal profiles around TSS regions for each processing combination from sample S05. The combination of bowtie2 with 5′-end trimming produces a distinct promoter-proximal signal profile resembling the divergence patterns observed in Figure 2C. (D) Mapping rates across processing combinations generated from different trimming and alignment strategies. Points represent individual samples, with lines connecting mean values and error bars indicating standard error of the mean.

Consistent with this clustering result, metaplot analysis showed that 5′-end trimming combined with bowtie2 produced a markedly altered TSS-centered signal profile, characterized by reduced promoter-proximal signal intensity relative to the other combinations (**Figure 3C**, representative sample; **Supplementary Figure S3D**, all samples). This profile closely resembled the PEPPRO-associated pattern observed in paired-end datasets (Figure 2C), linking the diagnostic workflow to the pipeline-level divergence identified above. The corresponding metaplot profiles remained altered when post-alignment filtering was omitted (**Supplementary Figure S3E**, representative sample; **Supplementary Figure S3F**, all samples). Representative locus-level examples showed the same divergence patterns at individual genes (**Supplementary Figure S3G**). Mapping-rate analysis showed that read retention remained relatively consistent across bwa-based combinations, whereas bowtie2-based combinations varied depending on trimming strategy (**Figure 3D**). Together, these observations indicate that structured divergence arises from interactions between processing steps, rather than from individual steps considered in isolation.

### Pipeline-defined assumptions constrain user-specifiable processing behavior

Having identified processing interactions associated with structured divergence, we next asked whether similar patterns would be observed in independently generated datasets under a shared library design. We generated paired-end PRO-seq libraries containing dual-end UMIs from human K562 cells and pig 3D4 cells, an alveolar macrophage cell line (see **Methods**). This design allowed us to evaluate pipeline behavior under a controlled library architecture while extending the analysis across species.

TSS-centered metaplot analysis showed that the three pipelines produced reproducible pipeline-associated signal profiles across both datasets (**Figure 4A**). In both human and pig PRO-seq libraries, PEPPRO showed reduced promoter-proximal signal intensity relative to proseq2.0 and nf-core/nascent under the shared dual-end UMI design. Locus-level tracks at orthologous TSEN15 loci further illustrated these patterns, with each pipeline producing characteristic coverage profiles in both species (**Figure 4B**; additional loci in **Supplementary Figure S4**). These observations indicate that the divergence pattern recurs across independently generated datasets and is not limited to the public K562 datasets analyzed above.

**Figure 4.**
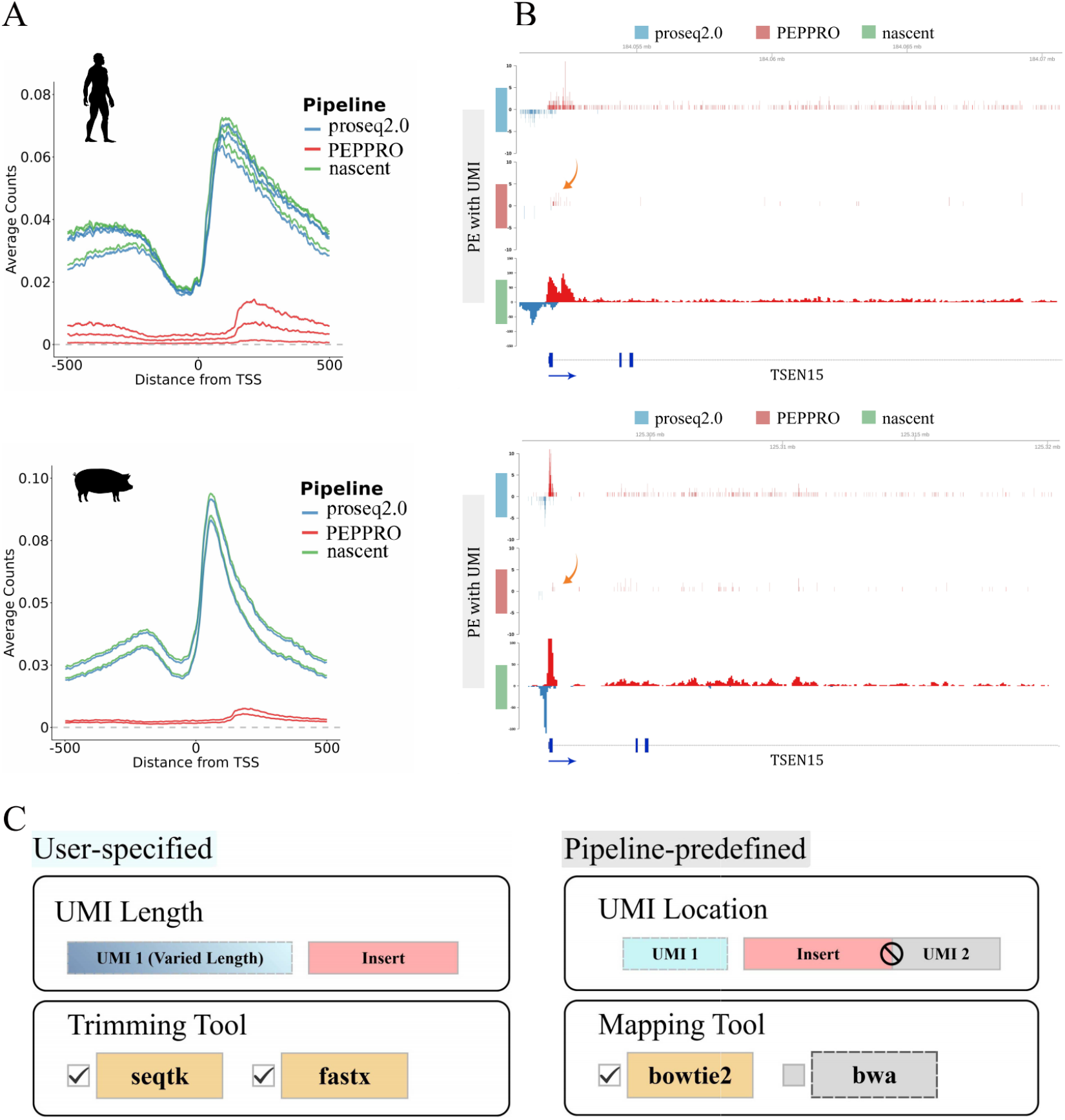
Pipeline-defined constraints shape reproducible divergence across species. (A) Metaplots of signal profiles around transcription start sites (TSS ±500 bp) for human and pig PRO-seq datasets generated under a shared library design (paired-end with UMIs), showing that similar divergence patterns are observed across species. (B) Genome browser tracks at orthologous TSEN15 loci in human (Sample S20) and pig (Sample S22) datasets, illustrating characteristic signal profiles associated with each pipeline under the same library design. (C) Schematic overview of user-specified parameters and pipeline-predefined assumptions in PEPPRO. User-specified settings include UMI length specification and trimming tool selection, whereas key assumptions such as UMI positional interpretation and supported aligner selection remain predefined by the pipeline. The pipeline assumes a fixed 5′ linkage between UMI and insert, limiting compatibility with dual-end UMI library architectures.

We next distinguished user-configurable parameters from predefined pipeline behavior. The extent of user control varied across pipelines. For example, PEPPRO allows users to select the preprocessing tool, specify UMI length and modify selected bowtie2 parameters (**Figure 4C**). Other aspects of processing, including aligner choice and the positional interpretation of UMIs relative to insert sequences, remain predefined. Consequently, the dual-end UMI library designs associated with the observed signal distortion cannot be fully accommodated through parameter adjustment alone. These observations indicate that compatibility depends not only on the parameters exposed to users, but also on how a pipeline interprets library structure.

### Library architecture and metadata availability shape reproducibility states across assays

To examine whether structured divergence extends beyond PRO-seq, we evaluated pipeline behavior across additional nascent RNA sequencing assays and metadata-availability contexts. We first analyzed published GRO-seq datasets from human BJ-5ta cells^49^, which use a paired-end library design without UMIs (**Supplementary Table S1**). Clustering based on TSS-proximal signal showed that samples grouped by analysis pipeline rather than by biological replicate (**Figure 5A**), consistent with the pipeline-associated divergence observed in PRO-seq datasets (**Figure 2A**). However, correlations between pipelines remained relatively high within the TSS-proximal signal window, indicating that regional signal intensity patterns were largely preserved.

**Figure 5.**
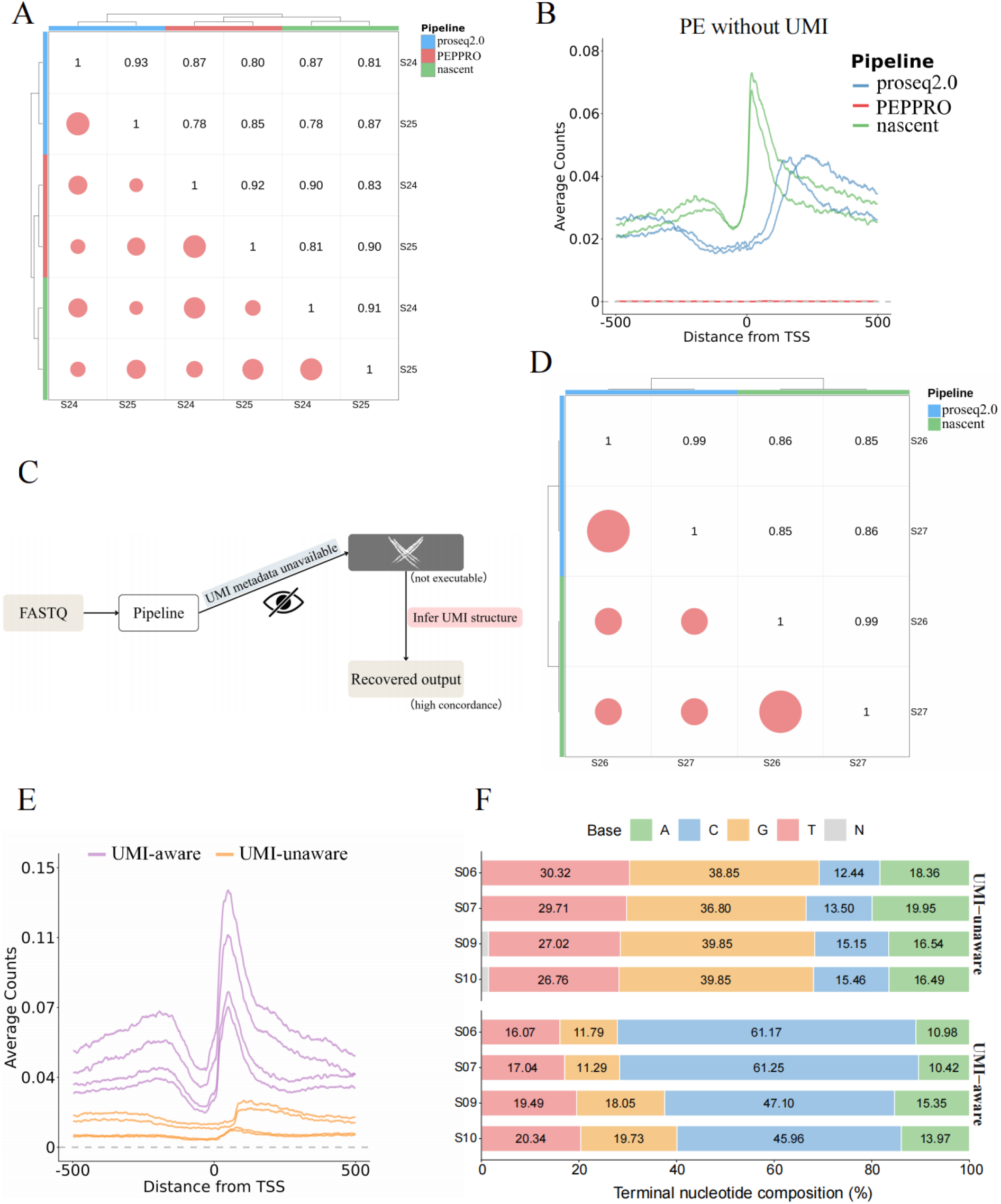
Assay-specific signal profiles and metadata availability shape reproducibility states. (A) Clustered heatmap of log_2_RPM-based correlations for GRO-seq data across TSS regions (±500 bp), showing that samples grouped by analysis pipeline rather than biological replicate despite relatively high overall correlations between pipelines. (B) Metaplots of GRO-seq signal profiles around TSS regions under a paired-end library design without UMIs, illustrating distinct positional signal profiles arising from pipeline-specific coordinate conventions. (C) Schematic illustration of PRO-cap processing under incomplete and recovered metadata conditions. Missing UMI-structure metadata prevents pipeline execution, whereas inference of hidden UMI architecture enables downstream processing. (D) Correlation clustering heatmap of PRO-cap samples processed using inferred UMI structures. Pearson’s correlation coefficients are shown in the upper triangle, and circle size in the lower triangle reflects correlation strength. (E) Comparison of UMI-unaware and UMI-aware processing for paired-end PRO-seq libraries with unreported terminal UMI structures. Retaining hidden UMI bases leads to reduced signal intensity, whereas removing inferred UMI sequences restores signal profiles. (F) Terminal nucleotide composition at inferred RNA 3′-end signal positions under UMI-unaware and UMI-aware processing. UMI-unaware processing retains terminal UMI bases at positions used to infer RNA 3′ ends, whereas UMI-aware processing removes inferred terminal UMIs before nucleotide composition is calculated. Stacked bars show the percentage of inferred RNA-strand A, C, G, T and N nucleotides for each sample.

Despite the high correlations, metaplot analysis of genome-wide signal tracks revealed distinct positional transcriptional profiles across pipelines (**Figure 5B**). This difference reflected distinct conventions for assigning transcriptional signal coordinates. nf-core/nascent reports signal using the 5′ end of reads, whereas proseq2.0 reports signal using the 3′ end. Near transcription start sites, 5′-end signals more closely mark transcription initiation positions, producing narrower and more upstream-enriched peaks. By contrast, 3′-end signals more closely mark RNA polymerase active-site positions and generate profiles resembling conventional promoter-proximal pausing patterns. Thus, the same GRO-seq libraries can produce distinct positional transcriptional profiles depending on how pipelines define signal coordinates.

PEPPRO showed an additional divergence pattern under this GRO-seq library design. Although PEPPRO remained relatively correlated with the other pipelines at the BAM-derived TSS-signal level (**Figure 5A**), BigWig-derived profiles showed markedly reduced signal intensity (**Figure 5B**; red curve). Pipeline inspection indicated a mismatch between the GRO-seq library orientation and the paired-end orientation assumptions used during PEPPRO alignment. This mismatch was associated with lower recovery of properly paired reads and altered strand-specific signal output (see **Methods**). These observations indicate that reproducibility at an upstream alignment-derived layer does not necessarily preserve downstream positional profiles. They further show that structured divergence can occur in paired-end assays even without UMI-related processing, placing the evaluated GRO-seq datasets within the distorted state.

We next analyzed PRO-cap datasets, a related nascent RNA sequencing assay with distinct library requirements. Because PEPPRO does not support PRO-cap data, this analysis used proseq2.0 and nf-core/nascent. Two publicly available K562 PRO-cap paired-end libraries were selected (**Supplementary Table S1**)^50^. Although the original study reported the presence of UMIs, detailed metadata describing UMI length and positional organization were not available. Under these metadata conditions, standard processing could not be executed because both pipelines required user-defined UMI length and position parameters (**Figure 5C**). Thus, these datasets initially occupied a non-executable state under the metadata available to downstream users.

We then asked whether the missing UMI structure could be inferred directly from the sequencing reads. Inspection of raw-read patterns indicated dual 6-bp UMIs, supported by terminal soft-clipping patterns and stepwise trimming analyses (see **Methods**). Incorporating the inferred UMI structure into the pipeline parameters enabled processing by both proseq2.0 and nf-core/nascent. TSS-proximal signal correlations were high across biological replicates and pipelines (**Figure 5D**): correlations were nearly identical between biological replicates (r = 0.99) and remained high between pipelines (r ≥ 0.85). TSS-centered profiles and genome-browser tracks also showed clear transcriptional signals after UMI-aware processing (**Supplementary Figure S5A–B**). These results indicate that the initial non-executable state arose from missing pipeline-relevant UMI metadata rather than from intrinsic incompatibility between the PRO-cap libraries and the pipelines.

This result prompted us to re-examine paired-end PRO-seq datasets for which UMI structures had not been reported. Inspection of raw-read patterns identified evidence of unreported terminal UMIs in four of the five datasets examined. We therefore compared UMI-unaware processing, in which the inferred UMI bases were retained, with UMI-aware processing, in which they were removed before downstream analysis. UMI-aware outputs showed higher concordance than UMI-unaware outputs (**Supplementary Figure S5C**). They also showed stronger promoter-proximal signal profiles characteristic of RNA polymerase pausing (**Figure 5E**; **Supplementary Figure S5D**).

Terminal nucleotide composition provided an additional readout of the effects of unreported UMIs. Under UMI-unaware processing, inferred RNA 3′-end positions were enriched for UMI-derived sequence features, with G and T as the dominant terminal bases in these libraries (see **Methods**). After removal of the inferred UMIs, the terminal profiles became cytosine enriched and resembled those of libraries without terminal UMIs (**Figure 5F**; **Supplementary Figure S5E**). This difference matters for interpretation because terminal nucleotide profiles are commonly related to RNA polymerase active-site positions, and cytosine enrichment has been associated with promoter-proximal pausing and elongation dynamics^22,27,51^. Thus, unreported terminal UMIs can alter both signal intensity and the inferred nucleotide context of active RNA polymerase. Analyses may therefore complete without error while assigning hidden UMI bases as RNA 3′ ends, resulting in silent signal loss and potentially misleading active-site profiles.

Finally, integrating observations across assays, library designs and metadata-availability conditions, we organized the evaluated datasets according to their empirical reproducibility outcomes (**Figure 6**). Single-end PRO-seq datasets produced consistent transcriptional signal profiles across pipelines and fell within the stable state. Paired-end PRO-seq and GRO-seq datasets exhibited structured divergence and occupied the distorted state. PRO-cap datasets processed under incomplete metadata conditions initially occupied the non-executable state, but transitioned to reproducible execution once essential UMI architecture was recovered. Paired-end PRO-seq datasets with unreported UMIs further illustrated how incomplete metadata can produce silent signal loss despite successful pipeline execution. Together, these results show that reproducibility states emerge from interactions among library design, pipeline assumptions and metadata availability, and that access to pipeline-relevant metadata is a critical determinant of reproducible transcriptional signal interpretation.

**Figure 6.**
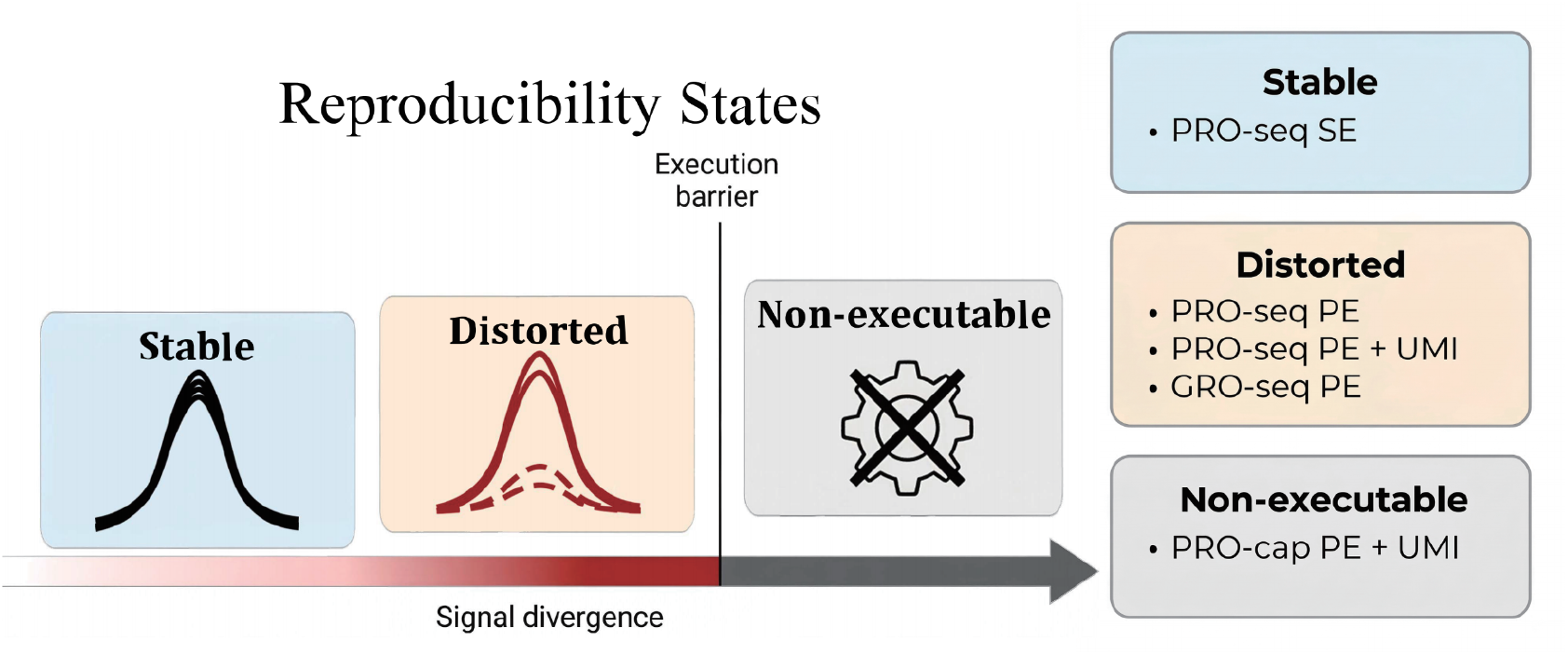
Reproducibility states defined by library design, pipeline assumptions and metadata availability. Conceptual summary of the reproducibility outcomes observed across the PRO-seq, GRO-seq and PRO-cap datasets evaluated in this study. Single-end PRO-seq datasets showed the stable state, whereas paired-end PRO-seq and GRO-seq datasets showed the distorted state. PRO-cap datasets under incomplete metadata conditions initially showed the non-executable state, but became executable after the required UMI architecture was inferred. Paired-end PRO-seq datasets with unreported terminal UMIs illustrate silent signal loss as a metadata-dependent outcome within otherwise executable analyses.

## Discussion

Computational reproducibility in sequencing analysis is often evaluated at the level of software versions, parameters and workflow execution. Our findings show that reproducibility also depends on how pipelines interpret library structure and convert sequencing reads into positional transcriptional signals. In the nascent RNA sequencing datasets evaluated here, these interactions produced distinct reproducibility states affecting signal recovery, RNA polymerase active-site interpretation and downstream quantitative estimates (**Figure 6**).

Within this framework, divergence is not simply a random consequence of tool variability. Instead, it reflects structured interactions between library design and pipeline assumptions. Previous studies of RNA-seq pipelines have shown that software choices and combinations of analytical components can substantially affect expression estimates and downstream predictions^52–54^. Our results extend this concern to nascent RNA sequencing, where assumptions governing trimming logic, mapping strategy, UMI organization, read orientation and signal-coordinate definition determine how sequencing reads are transformed into biologically interpretable positional profiles. Importantly, reproducibility can differ across analytical layers: pipelines may show similar upstream signal correlations while generating substantially different positional transcriptional profiles in downstream track-based analyses (**Figure 5A-B**).

Our results further show that sources of divergence differ in their accessibility to users. Library and protocol requirements define how sequencing reads should be interpreted. Some corresponding parameters, such as trimming-tool selection, UMI length and selected aligner settings, can be specified during pipeline execution. Other aspects, including aligner choice, UMI positional interpretation and orientation handling, may remain predefined within a pipeline (**Figure 4C**). Parameter flexibility therefore does not necessarily imply compatibility with all library architectures.

Metadata availability is therefore a central determinant of reproducible signal interpretation. This aligns with broader concerns that incomplete reporting of computational steps, software versions and parameter values limits reproducibility in RNA-seq analysis^2,8^. In nascent RNA sequencing, the relevant metadata extend beyond software settings to include pipeline-relevant library features, such as UMI length, position, read orientation and adapter structure. Missing or incomplete relevant metadata can make analyses non-executable when required parameters cannot be specified. A less obvious but particularly consequential outcome is silent signal loss, in which pipelines execute without error but biologically relevant transcriptional signals are reduced because hidden library features are not accounted for (**Figure 5E**). In the case of terminal UMIs, this loss extends beyond reduced signal intensity: unrecognized UMI bases can be mistaken for RNA 3′ ends, thereby distorting the inferred nucleotide context of active RNA polymerase. From this perspective, metadata availability is not only a documentation issue, but a prerequisite for making public datasets reusable by computational pipelines^2,31,32,55^.

Although our analysis focuses on nascent RNA sequencing, the underlying principle is likely relevant to other sequencing-based assays. Benchmarking studies in conventional RNA-seq, long-read transcriptomics, chromatin accessibility and chromatin conformation assays have shown that analytical choices and tool-specific assumptions can substantially alter downstream estimates or detected features^5,6,56–59^. These examples suggest that interactions between experimental design and pipeline assumptions are not unique to nascent RNA sequencing. Computational pipelines are often developed around specific experimental protocols, whereas library designs evolve through changes in read configuration, barcode structure, UMI placement or signal recovery strategy^29,30,35,60^. When these experimental changes are not matched by accessible pipeline-relevant metadata and clearly documented pipeline assumptions, reproducibility may fail at the level most relevant for biological interpretation. Consistent with this concern, pipeline-associated differences propagated to pausing-index estimates, illustrating how positional signal distortions can affect commonly used metrics of transcription regulation. For nascent RNA sequencing, this issue is especially consequential because positional signal recovery is not merely a visualization endpoint, but an input layer for downstream quantitative inference; distortions at this layer can therefore propagate into biological interpretations derived from read distributions^22–28^.

Together, these findings suggest that reproducibility should not be viewed as a binary property of computational pipelines. Instead, reproducibility states reflect whether biologically interpretable transcriptional signals can be consistently recovered under specific combinations of library design, pipeline assumptions and metadata availability. Organizing sequencing analyses according to stable, distorted and non-executable states provides a practical framework for interpreting analytical outcomes and anticipating incompatibilities between experimental designs and computational pipelines. More broadly, our results highlight the need to make pipeline-relevant metadata and pipeline assumptions explicit and accessible as sequencing protocols continue to diversify.

## Methods

### Published dataset collection

Publicly available nascent RNA sequencing datasets were collected from the Gene Expression Omnibus (GEO) and are listed in **Supplementary Table S1**. For the primary PRO-seq analysis, we selected 18 untreated human K562 datasets from 10 independent studies^16,19,28,35–41^. K562 datasets were prioritized to reduce biological heterogeneity and to enable comparison of pipeline behavior across different library designs within a shared cellular context. Library design annotations, including read configuration and the presence or absence of unique molecular identifiers (UMIs), were manually curated from published manuscripts, supplementary materials and GEO records. For cross-assay analyses, GRO-seq^49^ and PRO-cap^50^ datasets were obtained from GEO under accession numbers GSE113350 and GSE284853. Paired-end GRO-seq datasets without reported UMIs were included to evaluate divergence patterns in the absence of UMI-related processing. Dataset inclusion criteria included compatibility with the assays examined, availability of raw sequencing data and sufficient metadata to support pipeline execution or UMI-structure recovery analysis. UMI-containing libraries were prioritized where available.

### Cell culture and PRO-seq library preparation

K562 and 3D4 cells were cultured under standard conditions at 37°C with 5% CO₂. Both cell lines were maintained in RPMI-1640 medium supplemented with 10% fetal bovine serum (FBS) and 1% penicillin-streptomycin. Cells were used between passages 10 and 15 and were confirmed to be free of mycoplasma contamination.

PRO-seq libraries were prepared according to a previously described protocol with minor modifications^61^. Briefly, K562 and 3D4 cells were harvested on ice and washed with ice-cold PBS. For K562 cells, approximately 1 × 10⁷ cells per sample were permeabilized in cold permeabilization buffer and washed to remove endogenous nucleotides. For 3D4 cells, nuclei were isolated from approximately 1 × 10⁷ cells per sample under cold conditions using lysis buffer and used for the subsequent run-on reaction. Permeabilized K562 cells and isolated 3D4 nuclei were aliquoted for biotin run-on reactions.

Biotin run-on reactions were performed at 37°C in the presence of sarkosyl and four equimolar biotin-11-NTPs, each supplied at 1 mM, to label nascent transcripts associated with engaged RNA polymerase II. Run-on reactions were stopped by the addition of TRI Reagent LS. RNA was subsequently precipitated with absolute ethanol in the presence of GlycoBlue as a coprecipitant, and the resulting RNA pellet was washed with 75% ethanol. Purified RNA was fragmented by alkaline hydrolysis using 1 M NaOH, followed by neutralization with 1 M Tris-HCl, pH 6.8. Biotin-labeled nascent RNA was enriched using streptavidin magnetic beads and subjected to sequential library construction steps.

The enriched nascent RNA was ligated to a pre-adenylated 3′ RNA adapter, followed by a second round of streptavidin bead binding and stringent washing. The 5′ ends of RNA fragments were decapped using RNA 5′ pyrophosphohydrolase (RppH), repaired using T4 polynucleotide kinase (PNK), and ligated to a 5′ RNA adapter. The 3′ and 5′ RNA adapters contained 6-nt unique molecular identifiers (UMIs)^62^ to facilitate identification of PCR duplicates during downstream analysis. Adapter-ligated RNA was reverse transcribed, and cDNA was eluted from the beads before PCR amplification. Libraries were amplified using indexed primers for 10 PCR cycles and purified by bead-based size selection. The quality and concentration of final PRO-seq libraries were assessed before sequencing. Libraries were sequenced on an Illumina NovaSeq 6000 S4 platform using paired-end 150-bp reads.

### Pipeline execution and diagnostic workflow

Three nascent RNA sequencing analysis pipelines were evaluated: proseq2.0, PEPPRO and nf-core/nascent. All pipelines were executed using default configurations unless otherwise specified. Processing steps included adapter trimming, UMI handling, read alignment, post-alignment filtering and genome-wide signal track generation. Pipeline versions, execution parameters and software dependencies are listed in **Supplementary Table S2**.

To compare analytical strategies across pipelines, each pipeline was decomposed into key processing stages: UMI-aware trimming, read alignment, post-alignment filtering and signal generation. Differences in trimming strategy, mapping tool, filtering criteria, orientation handling and signal resolution were identified from pipeline documentation and verified using execution outputs and logs. For PEPPRO, user-specifiable settings included trimming tool selection, UMI length and selected bowtie2 parameters. Predefined pipeline assumptions included the use of bowtie2 as the mapping tool and default handling of UMI location and paired-end orientation. For GRO-seq analyses, orientation handling was examined to determine whether library orientation matched the paired-end orientation assumptions used during alignment.

To evaluate interactions between trimming and alignment strategies, we implemented a diagnostic workflow that systematically varied processing combinations using Snakemake^63^. Two trimming modes, dual-end trimming and 5′-end trimming, were combined with two aligners, bwa and bowtie2. Five paired-end PRO-seq datasets with UMIs were processed under each combination. After alignment, uniform post-alignment filtering was applied to all outputs, retaining properly paired reads that met the specified MAPQ > 10 threshold. Single-nucleotide signal tracks were generated from the filtered BAM files.

### Signal quantification and correlation analysis

Signal intensities were quantified from aligned BAM files as log_2_-transformed reads per million (log_2_(RPM)) within ±500 bp of annotated transcription start sites (TSSs). For gene body regions, defined as TSS + 2,000 bp to the transcription termination site (TTS), signals were quantified as log_2_-transformed transcripts per million (TPM). Pairwise Pearson correlation coefficients were calculated between samples and visualized by hierarchical clustering using the R package pheatmap (v1.0.13; https://cran.r-project.org/package=pheatmap) and ComplexHeatmap (v2.22.0)^64^. These analyses were used to assess similarity of transcriptional profiles across pipelines and diagnostic workflow conditions.

To prevent differences in native output-track resolution from confounding the comparisons, RPM and TPM values used for correlation analyses were calculated directly from aligned BAM files rather than from BigWig outputs. proseq2.0 and PEPPRO generate single-nucleotide-resolution tracks, whereas nf-core/nascent reports signal averaged over multi-base intervals by default. Datasets were stratified by library design as single-end, paired-end without reported UMIs or paired-end with UMIs. Within each group, pairwise Pearson correlation coefficients were calculated to assess the consistency of transcriptional profiles across samples and pipelines.

### Metaplot and locus-level visualization

Genome-wide signal tracks were aggregated around annotated TSSs within a ±500 bp window. To maintain consistent spatial resolution, single-nucleotide BedGraph files were used for nf-core/nascent, and BigWig files were used for proseq2.0 and PEPPRO. Average TSS-centered profiles were computed across genes using the R package genomation (v1.38.0)^65^. Metaplot analyses were used to compare positional transcriptional signal profiles across pipelines, diagnostic processing combinations and library designs.

Signal coverage tracks were visualized at individual genomic loci using the R package Gviz (v1.50.0)^66^. NELFB, ACTB, TSEN15, PRDX6, and QSOX1 were selected as representative expressed loci for visualization of promoter-proximal and gene-level signal patterns. Strand-specific signal distributions were examined across pipelines to assess consistency of transcriptional signal assignment.

### TSS-proximal heatmap visualization and pausing-index calculation

For TSS-proximal heatmap visualization, genes were ranked by the mean signal within the TSS-proximal region (±500 bp around the TSS), averaged across the three processing pipelines. The top 10% of genes were retained for plotting. The corresponding mean-signal cutoffs were 0.148 for paired-end libraries with UMIs, 0.156 for paired-end libraries without UMIs, and 0.381 for single-end libraries, resulting in 19,156, 19,303 and 19,154 selected TSS-centered windows, respectively. For each selected TSS-centered window, the region from 500 bp upstream to 500 bp downstream of the TSS was divided into 100 equally sized bins, and signal values were summarized within each bin. Heatmaps were generated separately for each library-design group and plotted using a common color scale across the three pipelines within the same group. For visualization, heatmap values were clipped to 0–1 for paired-end libraries and to 0–1.5 for single-end libraries.

Pausing index was calculated to assess whether pipeline-associated differences in promoter-proximal and gene-body signals propagated to downstream quantitative estimates. For each gene, promoter-proximal signal was quantified over the region from the TSS to 150 bp downstream of the TSS. Gene-body signal was quantified from 250 bp downstream of the TSS to the transcription termination site^67^. Signals in each region were normalized by the corresponding region length, and the pausing index was calculated as PI = (promoter-proximal region total read counts / region length) / (gene-body total read counts / region length). Non-finite values and values equal to zero were removed before log₂ transformation. Pausing-index distributions were visualized as log₂-transformed values.

### Mapping-rate, UMI inference and terminal nucleotide composition analysis

Mapping rates were calculated as the proportion of reads retained after alignment and filtering relative to total input reads. Mapping and filtering statistics were extracted from pipeline output files and workflow logs. Comparisons were performed across diagnostic processing combinations to assess variability in read retention.

To infer UMI presence and configuration in datasets lacking complete pipeline-relevant metadata, we implemented a read-structure inspection approach using raw sequencing reads. This procedure combined subset alignments, terminal soft-clipping inspection and stepwise terminal truncation. For PRO-cap datasets, alignment of a 5,000-read subset using bwa revealed abundant 6-bp soft clipping at both read termini. Stepwise trimming from either or both ends showed that removing 6 bp from both termini eliminated these soft-clipped bases, consistent with a dual 6-bp UMI architecture. The inferred UMI structure was then incorporated into downstream workflow parameters.

A similar inspection strategy was applied to paired-end PRO-seq datasets for which UMI structures were not reported in the available metadata. Datasets with evidence of unreported terminal UMI structures were processed under two conditions: UMI-unaware processing, in which terminal UMI bases were retained, and UMI-aware processing, in which inferred UMI sequences were removed before downstream analysis. Resulting signal profiles were compared to evaluate the effect of hidden UMI structures on signal recovery.

Terminal nucleotide composition was assessed from raw FASTQ reads to evaluate whether hidden terminal UMIs altered the inferred nucleotide context of RNA 3′ ends. Because these libraries capture nascent RNA in the reverse-complement orientation, RNA-strand terminal nucleotides were inferred from the corresponding read positions followed by base-complement conversion. For UMI-unaware processing, the read position that would otherwise be interpreted as the RNA 3′ end was extracted without accounting for UMI sequence. For UMI-aware processing, the terminal nucleotide was defined as the first base immediately downstream of the annotated or inferred UMI sequence. For paired-end libraries, this position was extracted from Read 1; for single-end libraries, the corresponding position was extracted from the single read. Extracted bases were converted to their complements (A to T, T to A, C to G and G to C), while N bases were retained as N. The proportions of inferred RNA-strand A, C, G, T and N nucleotides were calculated for each sample and visualized as stacked bar plots.

### Statistical analysis

All analyses and visualizations were performed using R v4.4.3 and Python v3.9.7. Statistical tests and multiple-testing adjustments are described in the corresponding figure legends where applicable.

## Data availability

All publicly available datasets analyzed in this study are listed in **Supplementary Table S1** together with their corresponding accession numbers. The newly generated PRO-seq data have been deposited in the Genome Sequence Archive (GSA) and Genome Sequence Archive for Human (GSA-Human) at the National Genomics Data Center (NGDC), China National Center for Bioinformation (CNCB). The human K562 PRO-seq data have been deposited in GSA-Human under accession HRA019677, whereas the pig 3D4 PRO-seq data have been deposited in GSA under accession CRA046448.

## Code availability

All scripts used for data processing, analysis, figure generation, UMI detection and parameter inference are available at https://github.com/xbzhou12138/Nascent-pipe.git. Pipeline execution configurations and parameter settings are provided in the repository and **Supplementary Table S2** to enable reproducibility of all analyses.

## Acknowledgements

We thank Daoyuan Wang, Xiaolong Qi, Shuheng Chan, and Yunxia Zhao for helpful discussions on library preparation. We are grateful to Prof. Shuhong Zhao for institutional guidance and support for this project. This work was supported by the Basic Research Project of Yazhouwan National Laboratory.

