## Supplemental Figures for "Hidden assumptions in nascent RNA sequencing pipelines define reproducibility states"

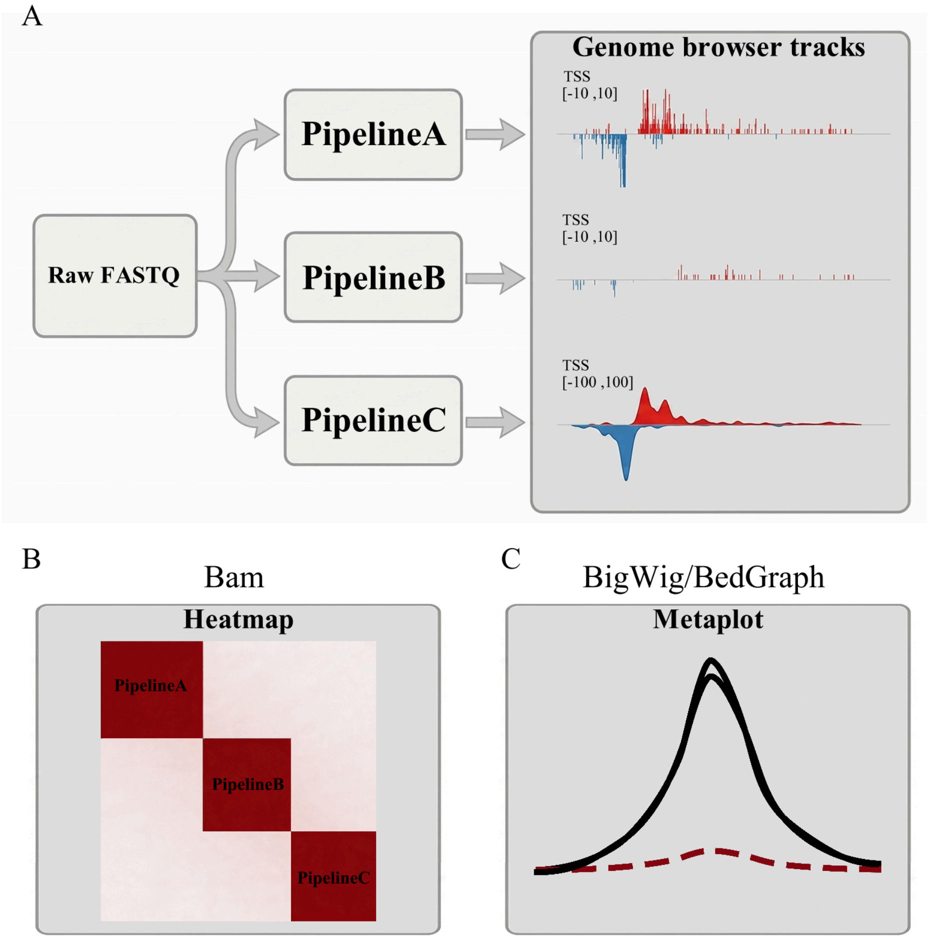


**Supplementary Figure S1. Overview of analytical outputs generated from identical raw FASTQ inputs processed using different nascent RNA sequencing pipelines.**

(A) Schematic representation of TSS-centered signal outputs generated by different pipelines, including differences in signal intensity and output resolution.
(B) Schematic representation of correlation clustering heatmaps used to compare signal profiles generated from aligned reads.
(C) Schematic representation of metaplots generated from genome-wide signal tracks produced by different pipelines.

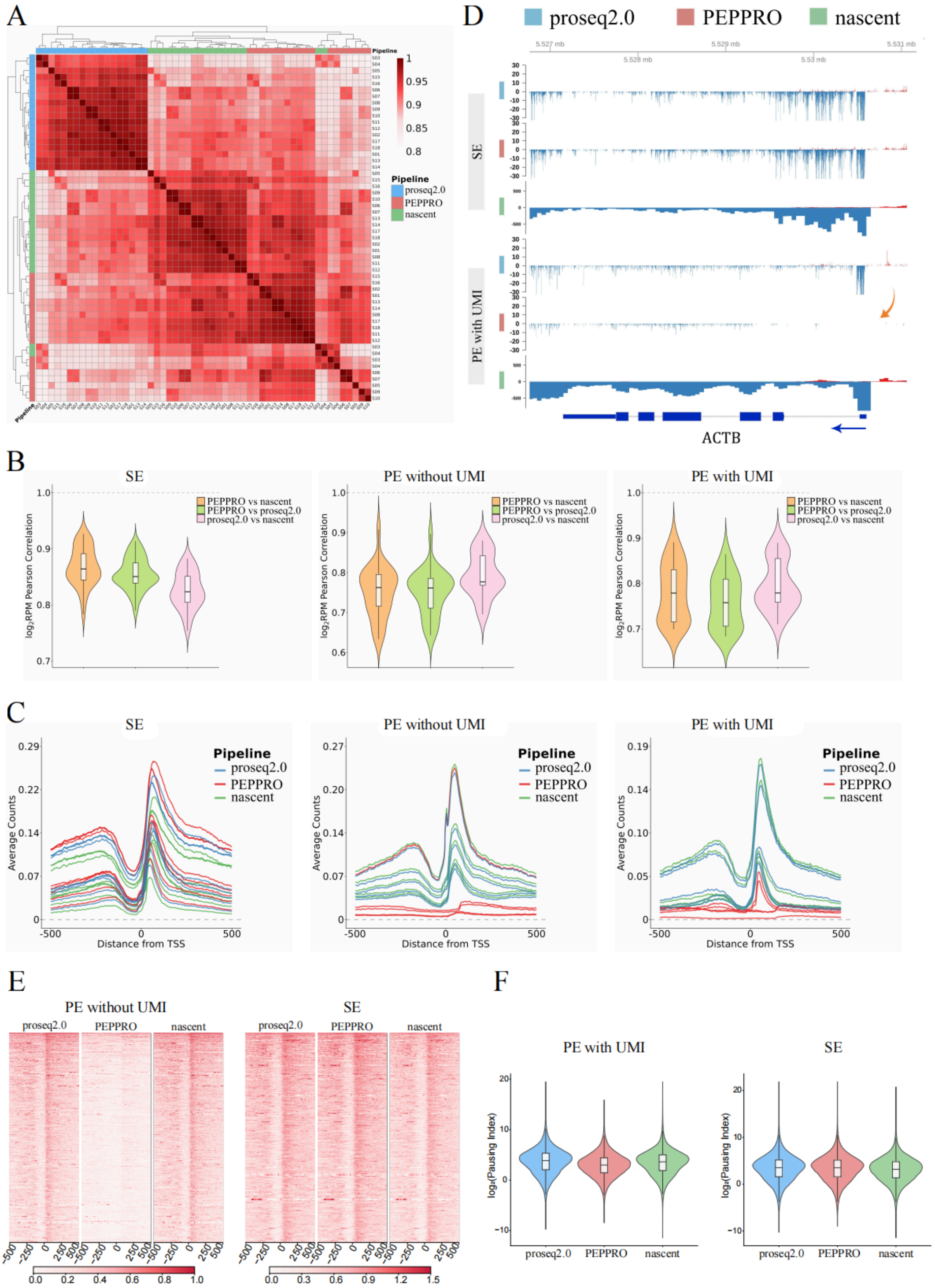

A
Pipeline
proseq2.0
PEPPRO
nascent
1
0.95
0.9
0.85
0.8
D
proseq2.0
PEPPRO
nascent
SE
PE with UMI
ACTB
B
SE
PE without UMI
PE with UMI
PEPPRO vs nascent
PEPPRO vs proseq2.0
proseq2.0 vs nascent
log2RPM Pearson Correlation
C
SE
PE without UMI
PE with UMI
Pipeline
proseq2.0
PEPPRO
nascent
Average Counts
Distance from TSS
E
PE without UMI
SE
proseq2.0
PEPPRO
nascent
F
PE with UMI
SE
log2(Pausing Index)
proseq2.0
PEPPRO
nascent

**Supplementary Figure S2. Divergence patterns across gene body regions and cross-pipeline comparisons.**

(A) Clustered heatmap of $log_2$TPM-based correlations for outputs from the three nascent RNA sequencing analysis pipelines across gene body regions, defined as TSS + 2,000 bp to TTS. The color gradient reflects correlation magnitude (red, high; white, low), and side color bars indicate the corresponding pipelines.
(B) Violin plots showing distributions of TSS-region correlations between pipelines within each library design group (single-end, paired-end without UMIs and paired-end with UMIs). The y-axis represents Pearson correlations of $log_2$RPM signals for pairwise cross-pipeline comparisons, with embedded boxplots indicating median and interquartile range. The dashed horizontal line marks a correlation coefficient of 1.
(C) Metaplots of TSS-centered signal profiles for the three library design groups. The x-axis indicates position relative to the TSS, and the y-axis shows average raw counts. Curves correspond to outputs from the three pipelines.
(D) Genome browser tracks at a representative locus (ACTB), illustrating local signal profiles for single-end (Sample S11) and paired-end with UMIs (Sample S01) libraries processed by each pipeline. The blue arrow indicates transcription direction and the pink arrow indicates signal loss; red and blue tracks denote signals on the positive and negative strands, respectively.
(E) Gene-level heatmaps of TSS-proximal signal intensities for paired-end libraries without UMIs (left, Sample S06) and single-end libraries (right, Sample S11). Rows represent TSS-centered windows from the top 10% of genes ranked by mean promoter-proximal signal across pipelines, and columns represent 100 binned positions across the TSS-centered region. Heatmaps are shown separately for proseq2.0, PEPPRO, and nf-core/nascent outputs. Heatmap values were clipped to 0–1 for paired-end libraries and to 0–1.5 for single-end libraries. Within each library-design group, a common color scale was used across the three pipelines.
(F) Violin plots showing distributions of $log_2$-transformed pausing index values across pipelines for paired-end libraries with UMIs and single-end libraries. Pausing index was calculated from length-normalized TSS-proximal and gene-body signal densities as described in **Methods**. Embedded boxplots indicate the median and interquartile range.

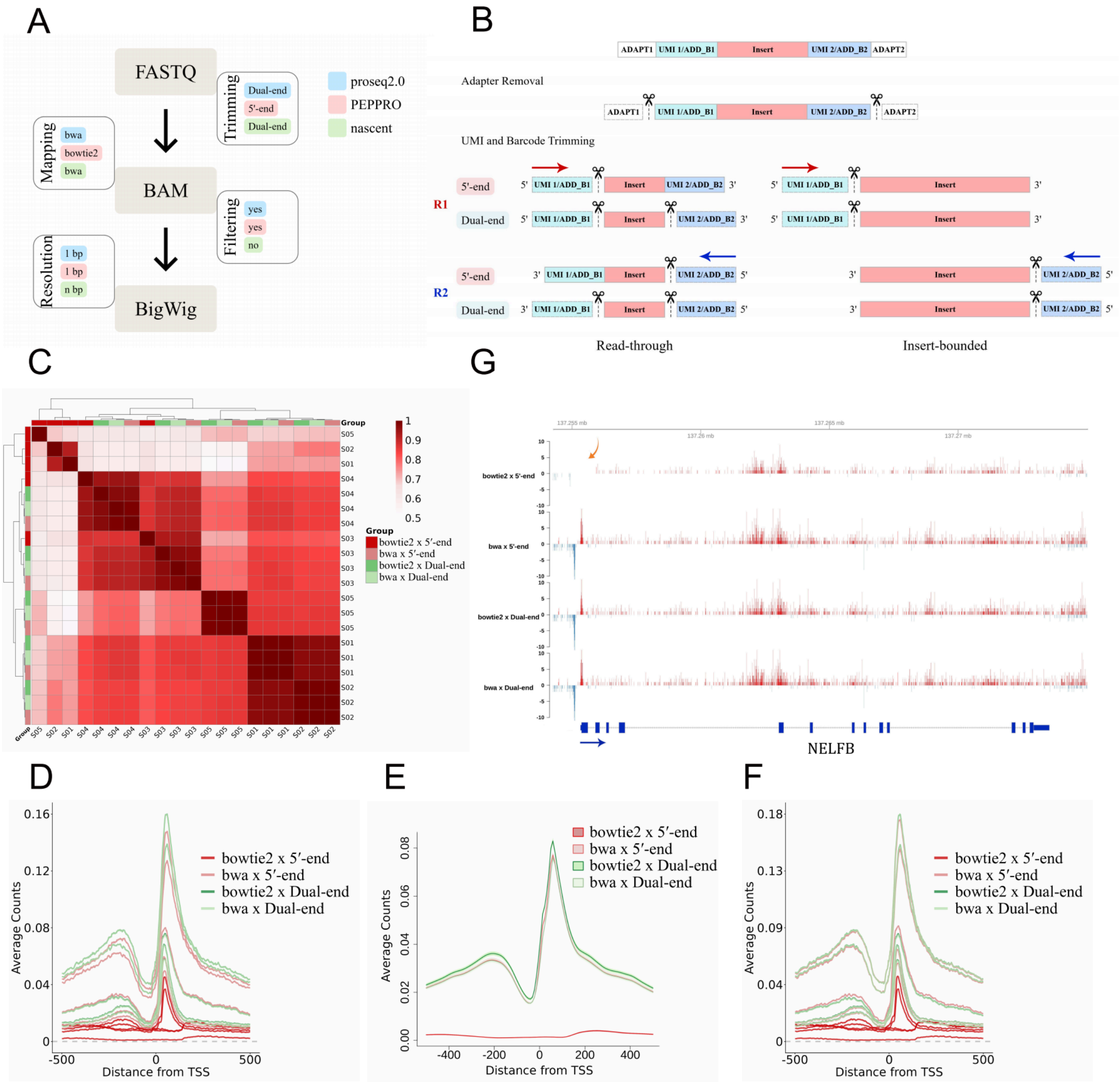

A
FASTQ
BAM
BigWig
Mapping
bwa
bowtie2
bwa
Trimming
Dual-end
5'-end
Dual-end
Filtering
yes
yes
no
Resolution
1 bp
1 bp
n bp
proseq2.0
PEPPRO
nascent
B
ADAPT1
UMI 1/ADD_B1
Insert
UMI 2/ADD_B2
ADAPT2
Adapter Removal
UMI and Barcode Trimming
R1
R2
5'-end
Dual-end
Read-through
Insert-bounded
C
Group
bowtie2 x 5′-end
bwa x 5′-end
bowtie2 x Dual-end
bwa x Dual-end
G
bowtie2 x 5'-end
bwa x 5'-end
bowtie2 x Dual-end
bwa x Dual-end
NELFB
D
E
F
Average Counts
Distance from TSS

**Supplementary Figure S3. Diagnostic workflow analysis of read trimming, alignment and post-alignment filtering.**

(A) Schematic overview of PRO-seq analysis, illustrating four processing stages (trimming, mapping, filtering and signal resolution) from raw FASTQ reads to BAM and BigWig signal files. Processing configurations implemented by the three analytical pipelines (proseq2.0, nf-core/nascent and PEPPRO) are indicated by distinct color blocks. In the mapping stage, nf-core/nascent can use different aligners but bwa is the default.
(B) Schematic illustration of different trimming strategies, showing how read-through status affects effective read boundaries under 5′-end and dual-end trimming.
(C) Clustered heatmap of $log_2$RPM-based correlations around TSS regions (±500 bp) without post-alignment filtering for processing combinations generated by the diagnostic workflow. The bowtie2 with 5′-end trimming combination forms a distinct cluster relative to the other combinations.
(D–F) Metaplots of signal profiles around TSS regions for diagnostic workflow conditions. (D) Metaplots generated from all samples after filtering. (E) Metaplot generated from sample S05 without filtering. (F) Metaplots generated from all samples without filtering.
(G) Genome browser tracks at the NELFB locus for sample S01, illustrating signal profiles for the four diagnostic processing combinations. The blue arrow indicates the direction of NELFB transcription and the pink arrow indicates the signal loss. Red and blue tracks denote signals mapped to the positive and negative strands, respectively.

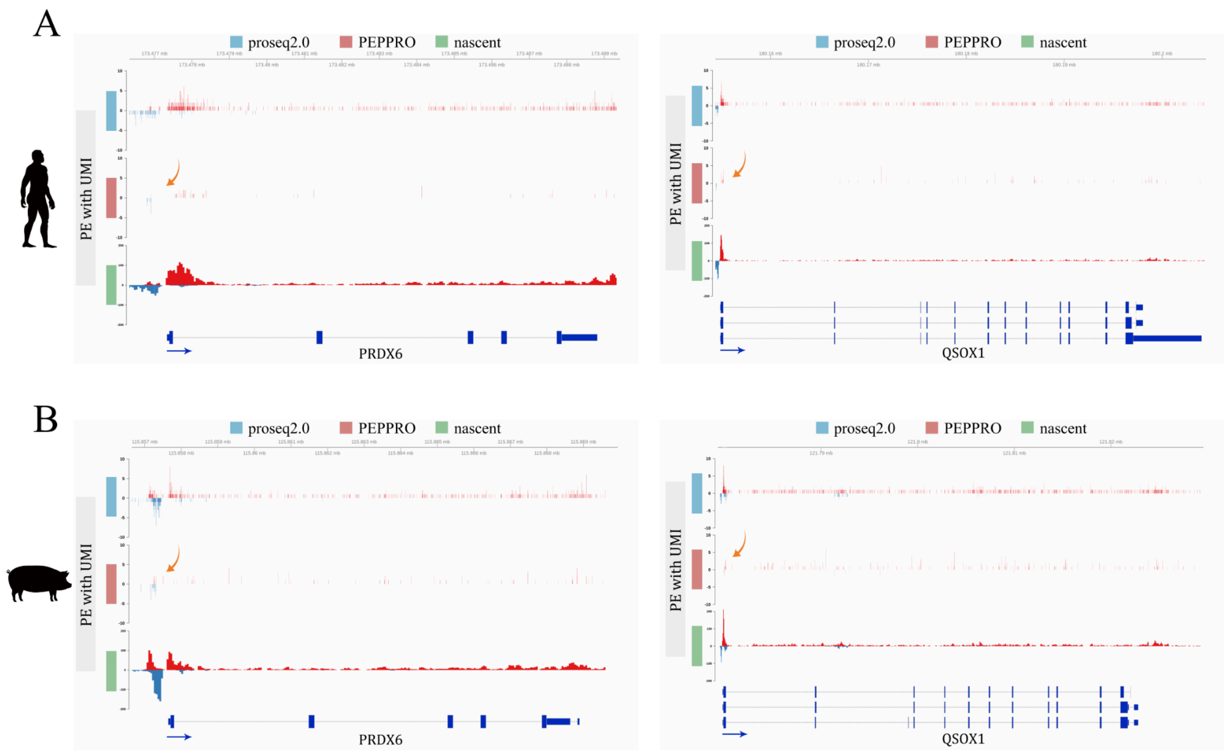


**Supplementary Figure S4. Genome browser tracks from independently generated human and pig PRO-seq libraries.**

(A) Genome browser tracks at the PRDX6 and QSOX1 loci from human sample S20.
(B) Genome browser tracks at the PRDX6 and QSOX1 loci from pig sample S22.

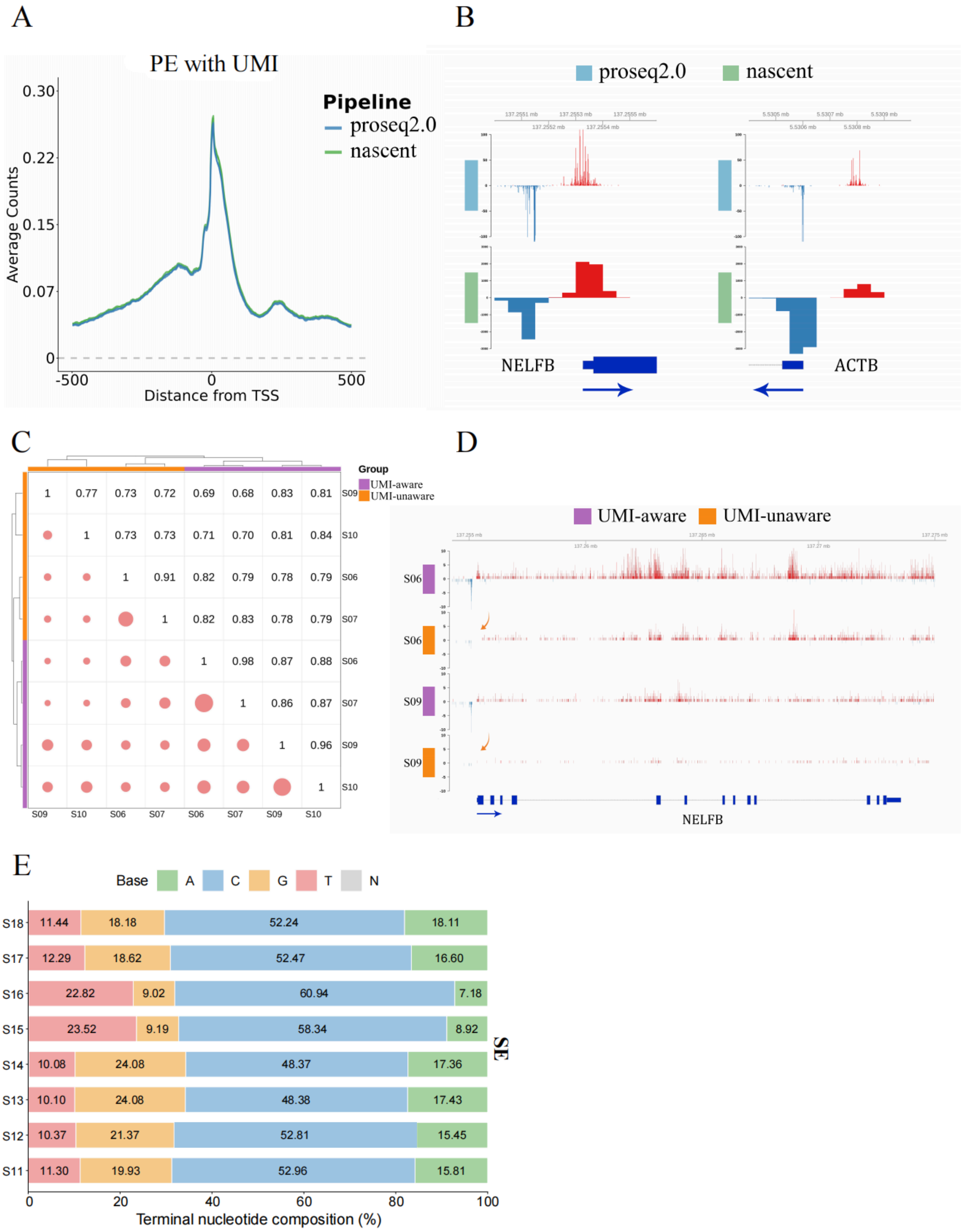
A
PE with UMI
Pipeline
proseq2.0
nascent
Average Counts
0.30
0.22
0.15
0.07
0
-500
0
500
Distance from TSS
B
proseq2.0
nascent
NELFB
ACTB
C
Group
UMI-aware
UMI-unaware
S09 1 0.77 0.73 0.72 0.69 0.68 0.83 0.81
S10 1 0.73 0.73 0.71 0.70 0.81 0.84
S06 1 0.91 0.82 0.79 0.78 0.79
S07 1 0.82 0.83 0.78 0.79
S06 1 0.98 0.87 0.88
S07 1 0.86 0.87
S09 1 0.96
S10 1
S09 S10 S06 S07 S06 S07 S09 S10
D
UMI-aware
UMI-unaware
S06
S06
S09
S09
NELFB
E
Base
A
C
G
T
N
S18 11.44 18.18 52.24 18.11
S17 12.29 18.62 52.47 16.60
S16 22.82 9.02 60.94 7.18
S15 23.52 9.19 58.34 8.92
S14 10.08 24.08 48.37 17.36
S13 10.10 24.08 48.38 17.43
S12 10.37 21.37 52.81 15.45
S11 11.30 19.93 52.96 15.81
SE
0
20
40
60
80
100
Terminal nucleotide composition (%)

**Supplementary Figure S5. Signal profiles and terminal nucleotide composition generated using inferred UMI structures in PRO-cap and PRO-seq libraries.**

(A) Metaplots of PRO-cap signal profiles after incorporation of inferred UMI structures into pipeline parameters, showing concordant profiles across pipelines and biological replicates.
(B) Genome browser tracks from PRO-cap sample S26 at representative loci after UMI-aware processing using inferred UMI structures.
(C) Correlation clustering heatmap of PRO-seq libraries grouped by UMI-unaware and UMI-aware processing conditions. Correlation coefficients are shown in the upper triangle, and circle size in the lower triangle reflects correlation strength.
(D) Genome browser tracks at the NELFB locus from samples S06 and S09, showing signal recovery under UMI-aware processing compared with UMI-unaware processing.
(E) Terminal nucleotide composition of recovered signal positions in single-end libraries. Stacked bars show the percentage of A, C, G, T, and N at terminal signal positions for each sample. Because these libraries do not contain terminal UMIs, they provide a non-UMI reference for comparison with the UMI-aware and UMI-unaware processing patterns shown in Figure 5F.
